# Acclimation capacity of two-banded seabream juveniles (*Diplodus vulgaris*) to port conditions: implications for the role of ports as nursery habitats

**DOI:** 10.1101/2025.10.04.680423

**Authors:** Céline M.O. Reisser, Matthias Rapp, Emilie Farcy, Eva Blondeau-Bidet, Florence Cornette, Kélig Mahé, Marc Bouchoucha

## Abstract

Ports are highly anthropized coastal environments characterized by elevated pollution and habitat alteration, posing significant challenges for marine organisms. However, the effects of port conditions on juvenile coastal fish using ports as nurseries remain poorly understood. Here, we combined field sampling and in situ caging experiments to investigate the genetic structure (RAD-seq), physiological condition (growth, lipid content, CYP1A-dependent biotransformation activity), and transcriptomic responses (RNA-seq) of juvenile two-banded sea bream (*Diplodus vulgaris*) exposed to port conditions. Juveniles from a port site were compared with individuals from two nearby non-port sites (T0), before fish from all populations were caged within the port for one month (T1). The site furthest from the port was slightly genetically differentiated. Port juveniles were larger and fatter at T0 than fish from non-port populations, and all populations showed differential regulation of lipid metabolism, xenobiotic detoxification, hypoxia response, and circadian rhythm pathways between T0 and T1. After one month of caging, weight differences persisted while muscle lipid content sharply declined in all populations. Only juveniles from the site furthest from the port significantly grew between T0 and T1. Port juveniles had a different transcriptional response to caging than non-port juveniles, with a higher expression of genes associated with energy metabolism, while non-port individuals overexpressed genes in detoxification, lipid metabolism, and oncogenic pathways. While we admit limitations related to caging conditions, our results suggest that port conditions strongly influence juvenile fish physiology and transcriptional levels, potentially promoting short-term acclimation strategies that could have long-term consequences for health and survival.

## Introduction

Port environments are among the most anthropogenically altered coastal habitats, characterized by intense pollution and extensive habitat modifications [1]. Industrialized ports receive a wide array of chemical contaminants, including heavy metals and organic compounds such as Polycyclic Aromatic Hydrocarbons (PAHs), Polychlorinated biphenyls (PCBs,) pesticides, and even pharmaceuticals [2–5]. Combined with physical habitat alterations such as replacement of natural substrates with artificial non-complex structures, dredging, artificial light at night (ALAN) or noise pollution, these stressors create challenging environments for marine organisms [6]. An estimated 50% of fish species depend on shallow coastal zones as nursery habitat [7,8], and the increased artificialization of the marine coastal environment degrades the quality of these habitats, reducing juvenile settlement, increasing juvenile mortality and/or disrupting adult population connectivity [9]. Moreover, port conditions can pose physiological, behavioral, and developmental risks for marine animals [10–15], particularly during early developmental stages [13,16].

Exposure to pollutants typically trigger a molecular response that aims at detoxifying the organism and may also trigger potential molecular alterations when defenses are overwhelmed. For instance, heavy metals can induce oxidative stress and disrupt metabolic and endocrine signaling, as reflected by altered expression of genes related to stress response, detoxification (*e.g.*, the cytochrome P450 family), and metal sequestration (metallothioneins) in exposed juvenile fish [17,18]. Endocrine-disrupting compounds such as nonylphenols and bisphenols further impair hormonal regulation [19]. Additionally, ALAN and underwater noise, both prevalent in port areas, have been shown to affect behavior and physiological health, with noise exposure specifically triggering oxidative stress, immune dysregulation, and neural responses in both fish and invertebrates [20].

To counteract the effects of long-term exposure to pollutants, resident species of heavily polluted ports (*i.e*. spending their entire life cycle in the port) can evolve genetic adaptations to tolerate these extreme conditions. A well-documented case is that of the Atlantic killifish (*Fundulus heteroclitus*), which inhabits PCB-contaminated estuaries and exhibits modifications in the aryl hydrocarbon receptor (AHR) pathway that reduce sensitivity to dioxin-like compounds, among which several Single Nucleotide Polymorphisms (SNPs) in coding regions [21,22]. These adaptations result from the presence of high selective pressures acting during the entire life of the organism, across multiple generations. In contrast, transient species that occupy ports only during specific life stages, like juveniles who use these areas as nurseries, would rather rely on short-term physiological acclimation to survive to high levels of pollution. Thus, despite lacking a genetic component to their response, transient species can also develop an adaptive response to pollution through physiological plasticity, by modulating molecular and enzymatic pathways involved in detoxification, cellular protection, and antioxidant defenses. For example, sticklebacks embryos (*Gasterosteus aculeatus*) that were experimentally exposed to copper showed reduced cellular stress response and lower disruption of ion homeostasis when later re-exposed, compared to fish that were not pre-exposed [23], illustrating a different regulation of molecular pathways as an adaptive response. However, to date, few studies have directly investigated adaptive responses of transient species to pollution directly within their natural environments.

Common garden experiments provide a powerful framework to test for the presence of history-dependent transcriptional response that could indicate acclimation, particularly when comparing genetically similar populations with contrasting histories of pollution exposure. In such a context, acclimation in individuals previously exposed to pollution could be reflected in reduced stress and detoxification responses, or more stable phenotypic and transcriptomic profiles, relative to naïve individuals. Fish originating from port environments are thus expected to exhibit enhanced tolerance to anthropogenic stressors, through pre-acclimated physiological states. By opposition, individuals from non-polluted populations may display stronger physiological and molecular stress responses after translocation. While common garden laboratory experiments are valuable for disentangling genetic and environmental effects, they often rely on simplified and standardized conditions that fail to capture the full complexity of natural environments, which is particularly the case for coastal marine ecosystems, and ports [24]. In contrast, *in situ* approaches that consists in translocating individuals directly into natural habitats exhibiting environmental conditions of interest, such as caging experiments, can better retain the environmental complexity, thereby providing more ecologically relevant insights into organismal responses to multiple, co-occurring stressors. In fish, common garden experiments through caging of individuals have been successfully employed to assess responses to pollutant exposure in port environments [11,25,26].

Here, we investigated the effects of port conditions and the presence of a putative acclimation to pollution in juveniles of the two-banded sea bream (*Diplodus vulgaris*) in the Bay of Toulon (north-western Mediterranean coast, France), a historically industrialized highly polluted port. We examined three sites (one originating from the port and two from nearby localities outside of the port) sampled at the end of their coastal growth phase, during which juveniles inhabit shallow nursery habitats for six to seven months before migrating offshore to join adult populations and analyzed them at two times: after sampling in their respective habitats (T0) and after one month of caging in port conditions (T1). We integrated physiological biomarkers with genomic and transcriptomic analyses to (i) verify the absence of significant genetic differentiation among populations, (ii) characterize the physiological and molecular responses that port conditions induce in the juveniles, and (iii) test for the presence of a possible adaptive response in fish juveniles that grew in the port, by comparing their response to caging to that of the non-port populations.

We hypothesized that (i) there should only be a small genetic differentiation across populations, (ii) the port conditions should result in differences in biometric characteristics and in levels of gene expression when comparing polluted and non-polluted sites at T0; (iii) the caging in the port should lead to a stable response of the port individuals between T0 and T1 and to a stronger response from non-port individuals (at least for key pathways related to detoxification); and (iv) at the end of caging, if an adaptive response was acquired by the port individuals, we should identify differences in biometric characteristics and gene expression at T1 between the port and the other two populations.

## Material and Methods

### Sampling

The study was conducted in the Bay of Toulon, located in the north-western Mediterranean Sea (Fig. 1). This semi-enclosed coastal system is composed of two distinct sub-basins: the Large Bay (42.2 km²) and the Small Bay (9.8 km²), separated by a 1.2 km breakwater constructed in the nineteenth century (Fig. 1). The Small Bay hosts major anthropogenic infrastructures, including one of the largest industrial harbors in France, the principal military harbor of the Mediterranean, and several marinas. Decades of naval, industrial, and urban activities have led to a severe multi-contamination of both sediments [27–30], the water column [29,31] and fish juveniles [3]. Notably, concentrations of mercury (Hg), lead (Pb) and copper (Cu) in sediments from the Toulon military harbor rank among the highest reported worldwide for marine environments [27]. In contrast, the Large Bay remains relatively less affected by direct human pressures [32–34]. This basin encompasses shallow rocky habitats that provide ecologically valuable nursery grounds for a wide range of juvenile fish species [35,36], thereby representing an important area for the maintenance of coastal biodiversity and fisheries productivity.

**Figure 1.**
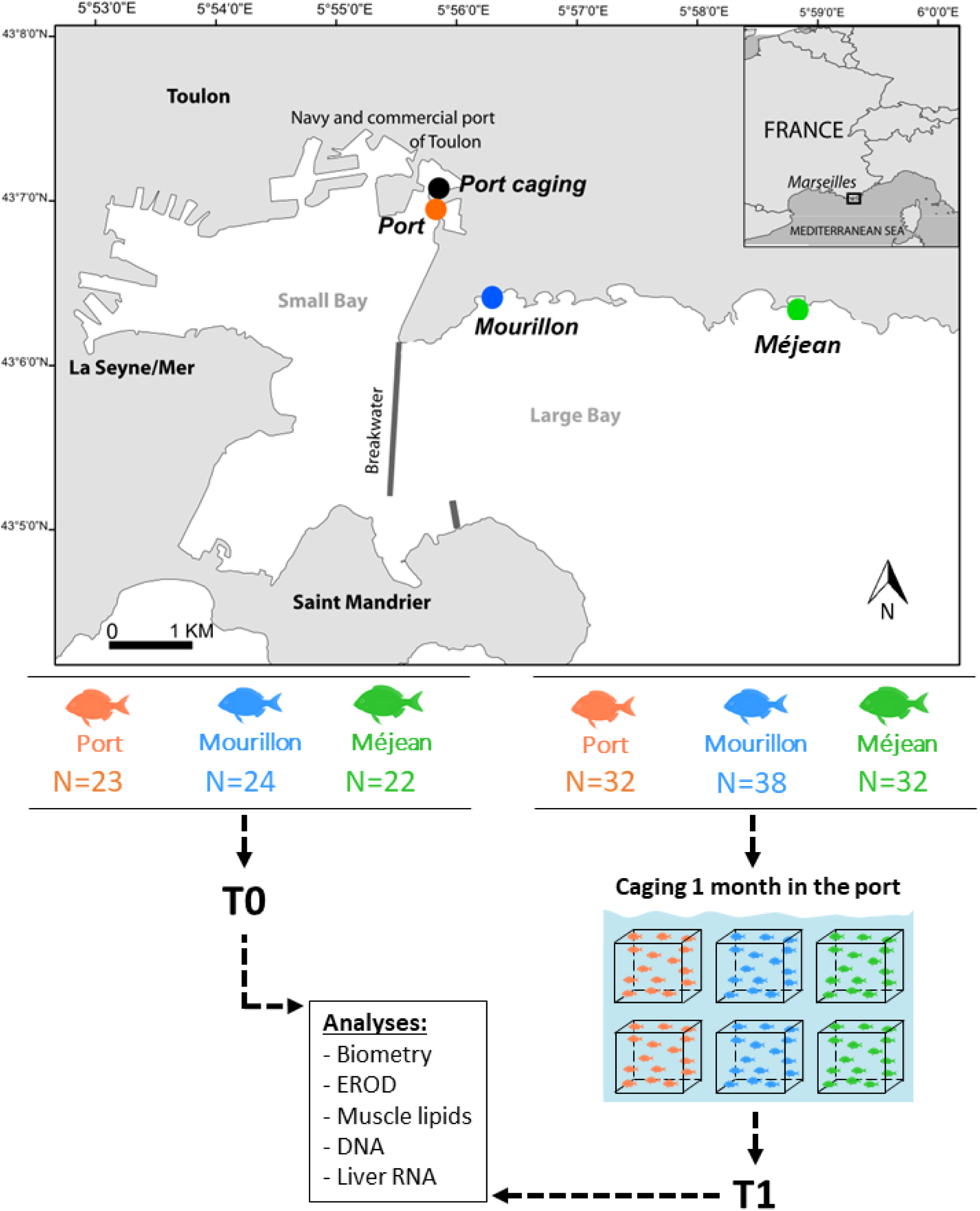
Sampling locations and experimental design of our study.

Three contrasted sampling sites were selected within the Bay of Toulon. The Toulon Port (43.112016, 5.929699) was chosen to represent the most polluted port environments present in the bay, while the other two sites (Mourillon Beach - 43.107415, 5.939506 - and Méjean Anse - 43.105844, 5.971684), located approximately 3 km and 5.5 km away from Toulon Port respectively, were selected as representative of more natural coastal habitats that serve as nursery grounds for juvenile rocky reef fishes.

During this study, a total of 171 *Diplodus vulgaris* fish of about six months of age were collected by scuba diving at night from three sampling locations (Port N=55, Mourillon N=62, and Méjean N=54, see Figure 1) between June 13^th^ and 15^th^ 2023. Sampling was performed as to standardize the size of the individuals sampled, minimizing the age difference. Sex was not recorded as the species is protandrous, with a sex maturation occurring after two years of age [37]. After capture, fish were transferred to 100 liters containers with continuous oxygenation while they were being processed. For each site, between 22 and 24 individuals per site (T0 samples) were directly euthanized through benzocaine overdose and liver, muscle, and caudal fin samples were collected, flash-frozen in liquid nitrogen before being stored at −80 °C until further processing (Figure 1). The remaining fish were placed in 0.6*0.6*1m cages (two cages per site containing between 16 and 19 fish) and immediately deployed within Toulon port (43.117425, 5.930431) over a duration of one month. To ensure feeding and reduce stress for the juveniles, small reefs made of oyster shells were placed in each of the cages, after having been kept in the port for one month prior to the experiment, to be colonized by epibionts and epiphytes that will feed the juveniles. After a month exposure (11^th^ July 2023), these fish (T1 samples) were euthanized and sampled following the same protocol described above (Figure 1).

### Biometric and physiological measurements

For each sampled fish, standard length (0.1 mm precision) and total weight (10 mg precision) were obtained. Muscle tissues from 120 fish (20 per location per time of sampling) were lyophilized over 72 hours prior to bead disruption, and sent to the platform LipidOcean, where ratios of triglycerides (TG), and esther sterols (ES) were determined. TG are the primary form of lipid storage in animal tissues, including fish muscle (high levels generally reflect greater energy reserves), while ES are membrane components that can give insight into cell membrane stability and physiological stress. For each condition, lipid analysis was performed with normalized per milligram of lyophilized tissue, and expressed in µg per milligram of dry weight.Difference in xenobiotic metabolic activity (CYP1A1) was examined in liver homogenates using Ethoxyresorufin-O-deethylase (EROD) activity. Specifically, liver sampled from 15 individuals from each population at T0, and from 20 individuals from T1 (10 individuals per cage at T1) were analyzed following the normalized protocol developed by AFNOR (NF T 90-385) [38]. Briefly, EROD activity was measured with a fluorometric assay of resorufin formation resulting from O-deethylation of the substrate, 7-ethoxyresorufin (Sigma/Merck, E3763) by CYP1A1 in presence of NADPH as a co-factor (Sigma/Merck, N6505). Kinetic measurements of the fluorescence intensity were made at an excitation wavelength of 532 nm and an emission wavelength of 587 nm, using black microtiter plates (Thermo Scientific), on a microplate reader TECAN® infinity. A five-point external calibration curve consisting of 0, 0.03125, 0.0625, 0.125, and 0.25µM of resorufin (Sigma/Merck, R3257) was used. Negative controls with no sample (blank) were included and the slope of the blank were subtracted from the slope of the samples. All activities were normalized by the amount of proteins contained in the samples, measured using the Bradford assay from Sigma-Aldrich (B6916) [39].

For all biometric and physiological measurements cited above, normality (*shapiro.test* function) in base R package “*stats*” and homoscedasticity (bptest function) from “*lmtest*” v0.9-40 R package [40] were evaluated. If the distribution followed normal rules, ANOVA was performed using a significance threshold of p<0.05 (anova function) in the “*stats*” package. Else, a non-parametric Kruskal-Wallis test was performed using the Kruskal.test function in the “*stats”* package. If significant differences were estimated, post-hoc Dunn test was performed (dunnTest function) from the R package “*FSA”* v0.10.1 [41], with the holm p-value adjustment method and significance threshold set at 5%.

### DNA extraction and RAD-seq bioinformatic pipeline

To test for the presence of genetic differences that could influence gene expression, gDNA was extracted from caudal fin tissue of 145 individuals (N=54, N=46 and N=45 for Port, Mourillon and Méjean respectively) with the Macherey Nagel Nucleospin Tissue kit following manufacturer’s recommendations, including the RNase treatment. gDNA quality and quantity was assessed with Nanodrop, Qubit, and 1% agarose gel migrations to verify compliance with quality requirements (Ratio 260/230 > 2; Ratio 260/280 > 1.8, no degradation) of the Montpellier Genomix platform (MGX, France; sample requirements: https://www.mgx.cnrs.fr/index.php/mgx/description-des-plateaux/mgx-ngs-igf). Samples were standardized at a concentration of 40ng.µL^-1^ and 2µg of DNA per sample were sent to MGX for RAD-seq library preparation using the sbfI restriction enzyme. Libraries were sequenced on an Illumina Novaseq SP1 FC PE150 and then demultiplexed by the platform.

Quality of raw reads was assessed using FastQC v0.12.1 (https://github.com/s-andrews/FastQC). Raw sequences were filtered using fastp v1.3.2 [42]: specifically reads that showed an average quality less than 28 and a length below 50 were removed. We also used fastp to remove adapter sequences, and poly-A tails that are typical of Illumina Novaseq sequencers. Filtered reads were mapped onto the *Diplodus sargus* reference genome (https://www.ncbi.nlm.nih.gov/assembly/GCA_903131615.1/) using BWA-mem v2.3 [43] and default settings. The resulting mapped files were filtered using Sambamba v1.0.1 [44], and reads that were not correctly paired and were not primary alignment were removed. We used Stacks v2.68 [45] and the ref_map.pl script to call for the genotypes, and required SNPs to be present in all three sites and loci to have less than 20% missing data (-X “populations: -p 3 -r 0.8”) and also removed loci that were in Hardy Weinberg disequilibrium (-hwe parameter). Stacks originally reconstructed 1648588 loci with an effective coverage per sample of 37.5X and identified 7793992 Single Nucleotide Polymophism (SNP) sites. The filtration parameters led to the conservation of 49666 loci and 1286716 SNPs. We then used BCFtools v1.23 [46] to remove loci that had a depth less than 15 (763559 SNPs kept), and had a minor allele frequency lower than 0.05 (181599 SNPs kept). VCFtools v0.1.16 [47] was used to estimate individual heterozygosity, average loci depth and percentage of missing genotypes, with eight individuals removed which had more than 25 % missing genotypes.

### Population genetic structuring and differentiation

To visualize the relationship among individuals and sites, we performed a Principal Component Analysis (PCA) on the individuals’ multigenotypes using the dudi.pca function of the R package “*adegenet*” v2.1.11 [48], keeping 4 PC axes. We performed unsupervised clustering with the find.clusters function, and assessed the best number of groups using the smallest Akaike’s information criterion (AIC). We calculated pairwise site F_ST_ using the stamppFst function of the “*StAMPP”* v1.6.3 package [49], with 10000 bootstrap and assessed significance using a confidence interval of 95%.

### RNA extraction and RNA-seq bioinformatic pipeline

mRNA was extracted from frozen liver samples (7 samples per site and per sampling time, resulting in 42 samples in total) using the Macherey Nagel Nucleospin RNA kit (Düren, Germany), following manufacturer’s recommendations. RNA quantity and purity was assessed using a Nanodrop. RNA quality was evaluated on a Bioanalyzer, and 200ng of RNA per sample with RIN>7 was sent to the Montpellier Genomix platform (MGX, France; sample requirements: https://www.mgx.cnrs.fr/index.php/mgx/description-des-plateaux/mgx-ngs-igf), for library preparation and sequencing. Libraries were constructed using the TruSeq kits and sequenced on an Illumina Novaseq S4 chip.

Quality of raw reads was assessed using FastQC v0.12.1. Raw sequences were filtered using fastp v1.3.2: specifically reads that showed an average quality less than 28 and a length below 50 were removed, to ensure efficient downstream analyses. We also used fastp to remove adapter sequences, and poly-A tails that are typical of Illumina Novaseq sequencers. Filtered reads were mapped onto the sister species’ *Diplodus sargus* reference genome (https://www.ncbi.nlm.nih.gov/assembly/GCA_903131615.1/) using star v2.7.10b [50] and default settings. The resulting mapped files were sorted using Sambamba v1.0.1. Finally, we used the conda package subread v2.0.8 and the FeatureCounts command [51] with default parameters to obtain the raw read count table per gene for each sample. The table was further processed using “*DESeq2*” v1.50.2 R package [52] to remove genes detected in fewer than two individuals (out of all individuals) and with a read count below 0.5 counts per million (CPM), resulting in a matrix of 21395 genes and 42 samples.

### Redundancy Analysis on global gene expression data

To observe the variations in individuals’ read count in the global dataset (all the 21395 genes of our dataset), and assess the contribution of the population, the pollution background, the time of sampling, and their interaction, we first applied a variance stabilizing transformation on the raw read counts using the vst function of “*DESeq2”* R package. We then performed a Redundant Analysis (RDA) on the transformed expression table using the rda function of the “*vegan”* v2.7-2 R package [53]. We tested the model “*expression ∼ population * time + pollution * time”* with ANOVAs (anova function in the “*stats*” package), with 999 permutations and a significance threshold set at p-value=0.05.

We also tested the model “*expression ∼ population + pollution*” to assess the contribution of the population and pollution background within each sampling time (one ANOVA for T0 samples, and one ANOVA for T1 samples), with 999 permutations and a significance threshold set at p-value=0.05.

### Differential Expression Analysis (DEA)

All DEA were performed on the raw count matrix (21395 genes). First, to study the impact of port environmental conditions, we performed a DEA by comparing individuals originating from Port to those originating from Méjean and Mourillon (setting the reference as the less impacted population for each comparison), using “*DESeq2*” and the variable “population” as the contrast. Genes were considered significantly differentially expressed if p-value < 0.05 with a false discovery rate (FDR) of 5 %. GO term enrichment was performed in Reactome [54] using the gene symbol of the enriched genes, a significance at p-value<0.05 with a false discovery rate (FDR) of 5 %. In addition, KEGG pathway enrichment analysis was performed on the DEGs using the *Larimichthys crocea* database in KOBAS-i [55], with significance defined as p-value<0.05.

Second, to compare the specific response of each population to one month of caging in polluted conditions and test for the presence of acclimation, we first performed DEA between T0 and T1 samples within each of the three populations. The “time” variable was set as contrast with T0 as a reference, and statistical significance as well as GO terms and KEGG pathways enrichment were performed for each DEA as explained above.

Finally, to test if the Méjean and Mourillon fish eventually reached a similar transcriptomic state as Port individuals once exposed to port conditions, or if differences remained after a month (possibly indicating an adaptive response), we conducted a DEA and a functional enrichment analysis on the T1 samples, with the same set up as for T0.

## Results

### Biometric and physiological measurements

#### T0 results

At T0, individuals from Port were significantly longer (ANOVA p-value=2.91E-05) and heavier (Kruskall-Wallis p-value=1.74E-04) than those of Méjean and Mourillon (mean=55.48 ± 6.16mm, 48.24 ± 4.02mm and 51.01 ± 3.93mm, and mean=2.49 ± 0.83g, 1.55 ± 0.42g and 1.83 ±0.45g respectively; Supplementary Table 1; Table 1; Figure 2A and B).

**Figure 2.**
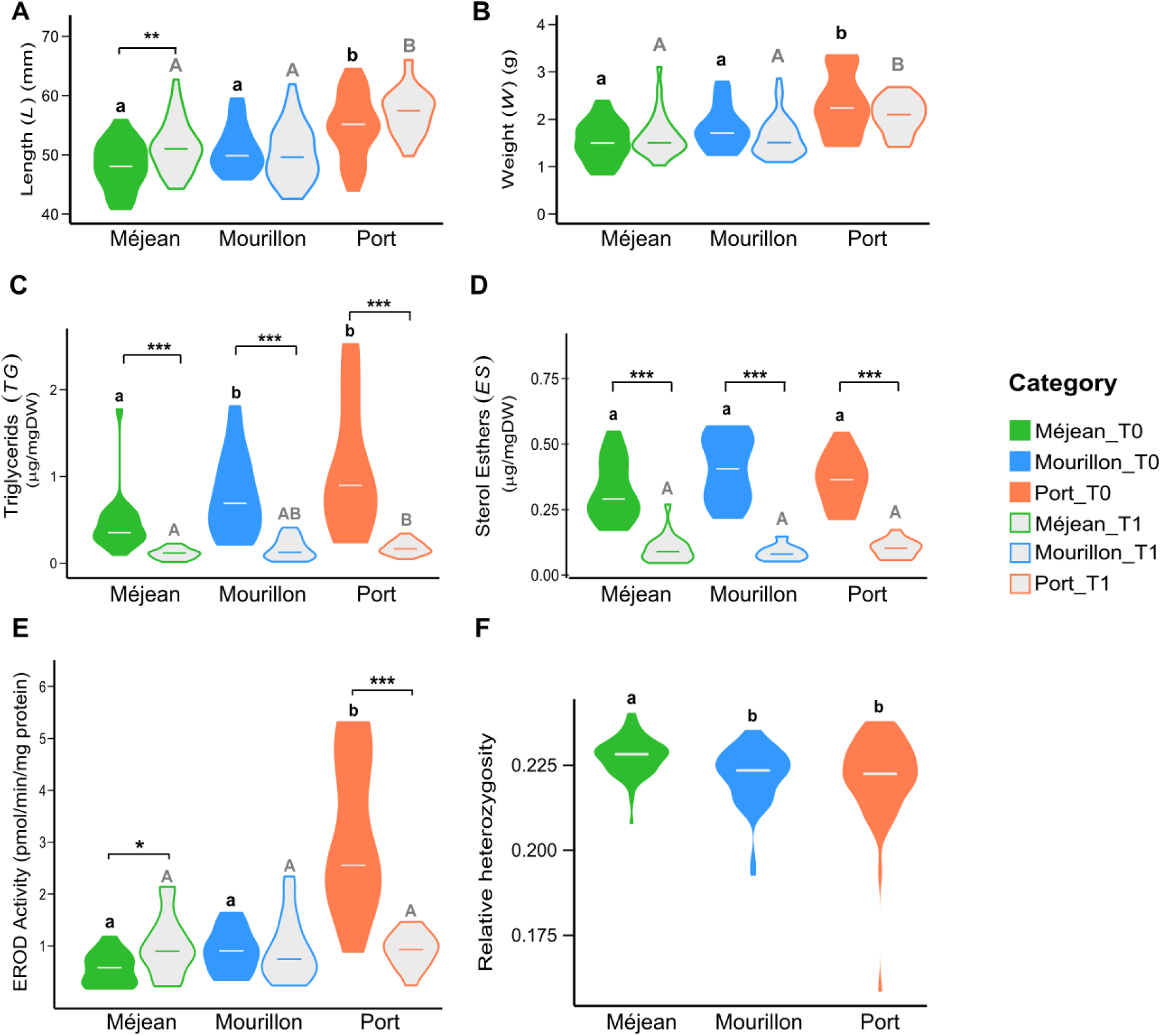
Biometry, physiology and heterozygosity results: (A) total length, (B) weight, (C) triglycerides muscle content, (D) ester sterols muscle content, (E) liver EROD activity and (F) individual’s relative heterozygosity. Letters show the results of the statistical tests (AMOVA or Kruskal-Wallis; small caps within T0, large caps within T1). Similar letters indicate no significant difference. Stars indicate the result of the statistical test between T0 and T1 within each population (*: p-value<0.05; **: p-value< 0.01; ***: p-value<0.001).

**Table 1.** Biometric statistical analysis results. Bold indicates significance at 0.05.

| Parameter | Time | Test | p-value |
| --- | --- | --- | --- |
| Length | T0 | ANOVA | **2.91E-05** |
|  |  | Méjean_vs_Mourillon | 0.078 |
|  |  | Port_vs_Méjean | **1.20E-04** |
|  |  | Port_vs_Mourillon | **0.044** |
|  | T1 | ANOVA | **1.51E-09** |
|  |  | Méjean_vs_Mourillon | 0.367 |
|  |  | Port_vs_Méjean | **7.39E-08** |
|  |  | Port_vs_Mourillon | **1.27E-05** |
| Weight | T0 | Kruskal-Wallis | **1.74E-04** |
|  |  | Méjean_vs_Mourillon | 0.105 |
|  |  | Port_vs_Méjean | **1.03E-04** |
|  |  | Port_vs_Mourillon | **2.60E-02** |
|  | T1 | Kruskal-Wallis | **1.02E-05** |
|  |  | Méjean_vs_Mourillon | 0.794 |
|  |  | Port_vs_Méjean | **1.26E-04** |
|  |  | Port_vs_Mourillon | **3.41E-05** |
| EROD | T0 | Kruskal-Wallis | **1.51E-06** |
|  |  | Méjean_vs_Mourillon | 0.079 |
|  |  | Port_vs_Méjean | **9.10E-07** |
|  |  | Port_vs_Mourillon | **1.64E-03** |
|  | T1 | Kruskal-Wallis | 0.796 |
| TG | T0 | Kruskal-Wallis | **1.61E-03** |
|  |  | Méjean_vs_Mourillon | **2.10E-02** |
|  |  | Port_vs_Méjean | **2.04E-03** |
|  |  | Port_vs_Mourillon | 0.298 |
|  | T1 | Kruskal-Wallis | **4.70E-02** |
|  |  | Méjean_vs_Mourillon | 0.302 |
|  |  | Port_vs_Méjean | **4.20E-02** |
|  |  | Port_vs_Mourillon | 0.298 |
| ES | T0 | Kruskal-Wallis | 0.069 |
|  | T1 | Kruskal-Wallis | 0.282 |

Levels of TG differed among populations (Kruskall-Wallis p-value=1.61E-03) with Méjean showing the lowest level (mean=0.46 ± 0.34 µg.mg(DW)^-1^) while Mourillon and Port had statistically similar levels (mean=0.88 ± 0.62 µg.mg(DW)^-1^ and mean=2.10 ± 1.87 µg.mg(DW)^-1^ respectively; Figure 2C; Supplementary Table 2; Table1). Levels of ES were similar across all sites at T0 (Figure 2D; Supplementary Table 2; Table 1).

EROD activity showed a strong spatial gradient (Supplementary Table 3), with an activity in Port being statistically higher (Kruskall-Wallis p-value=1.51E-06) than that of Méjean and Mourillon (mean=4.19 ± 3.13 pmol.min^-1^.mg(prot)^-1^, 0.56 ± 0.31 pmol.min^-1^.mg(prot)^-1^ and 1.03 ± 0.58 pmol.min^-1^.mg(prot)^-1^ respectively; Table 1; Figure 2E).

#### T1 results

At T1, differences in length and weight among sites persisted (ANOVA p-value=1.51E-09 and Kruskal-Wallis p-value=1.02E-05 respectively), with Port individuals remaining bigger and heavier (mean=57.27 ±3.77mm) than those of Méjean and Mourillon (Table 1, mean=51.48 ± 4.28mm and 50.90 ± 6.08mm respectively; Figure 2A and B).

Levels of TG differed between Port and Méjean (mean=0.19 ± 0.10 µg.mg(DW)^-1^ and 0.11 ± 0.06 µg.mg(DW)^-1^ respectively, p-value=1.61E-03), but were not different between Méjean and Mourillon or between Port and Mourillon (Supplementary Table 2; Table 1, Figure 2C). Levels of ES did not differ among individuals from the three sites after a month of caging at T1 (Figure 2D).

Also, EROD activity did not significantly differ among individuals from the three sites after one month of caging (Table 1, Figure 2E).

#### T0 versus T1 results

Intra-site comparisons (T0 versus T1) revealed that the fish from all three populations did not significantly increase in weight, and only Méjean individuals significantly increased in length (ANOVA p-value=0.006; Figure 2A and B).

Both TG and ES showed changed between T0 and T1 in all populations, with higher lipid reserves at T0, followed by a marked decrease at T1 (TG: p-value=2.61E-06, p-value=4.23E-07, p-value=1.55E-06 and ES: p-value=6.30E-068, p-value=2.38E-07, p-value=6.30E-08 for Port, Méjean and Mourillon respectively; Figure 2C and D).

EROD activity significantly increased in Méjean (mean=1.18 ± 0.80pmol.min^-1^.mg(prot)^-1^ ; Kruskal-Wallis p-value=0.013) and decreased in Port (mean=1.62 ±1.71pmol.min^-1^.mg(prot)^-1^ ; Kruskal-Wallis p-value=3.09E-05) while remaining unchanged in Mourillon (mean=1.23 ± 1.06pmol.min^-1^.mg(prot)^-1^ ; Figure 2E).

### Population genetic diversity and structuring

Observed heterozygosity among sites (Figure 2F) showed that the Méjean individuals had a higher genetic diversity compared to Port and Mourillon populations (Kruskal-Wallis p-value= 0.001; Dunn Test p-value=0.005 and 0.002 respectively), while Mourillon was not different from Port. PCA analysis did not show the presence of genetic structuring, both according to time of sampling and to geographical origin (Figure 3A). Unsupervised hierarchical clustering failed to identify more than one genetic cluster. Pairwise FST values were small and only significant when Méjean was involved in the comparisons (FST=8.80E-05, p-value=0.036 for Méjean vs. Port; FST=2.09E-04, p-value=1.00E-04 for Méjean vs. Mourillon; FST=3.16E-05, p-value=0.252 for Mourillon vs. Port).

**Figure 3.**
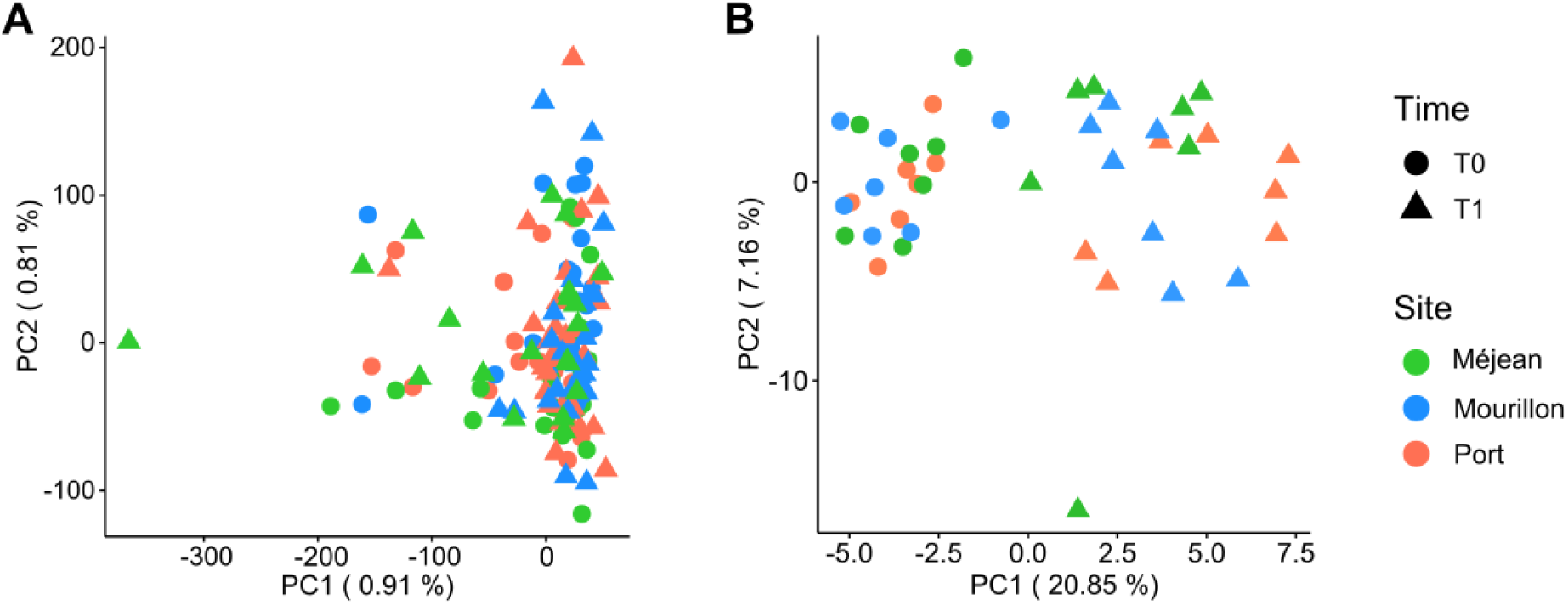
Multivariate analyses. (A) Principal Component Analysis of individuals’ genotypes and (B) Redundancy Analysis of individuals’ log transformed read count (vst transformation), according to the site of origin and the time of sampling.

### Gene expression analysis

#### RDA on expression profiles

On the global dataset (including both T0 and T1 samples), our model was significant in explaining variation in expression across samples (R^2^adj=0.18; ANOVA p-value<0.001) and showed a strong effect of the time of sampling (ANOVA p-value<0.001), while the population of origin did not significantly contribute to the global variance (ANOVA p-value=0.132). We also found that the pollution background of the individuals had a weak contribution (ANOVA p-value=0.063) (Figure 3B).

Within the T0 samples, the model was not significant. However, within the T1 samples, both the pollution background and the population of origin had a significant impact (ANOVA p-value=0.022 and 0.014 respectively), with samples seemingly segregating into three clusters, with Mourillon as the intermediate one (Figure 3B).

#### DEA and functional enrichment at T0

DEA identified 23 differentially expressed genes (DEGs) between Port and Méjean (9 downregulated, 14 upregulated), and 63 DEGs between Port and Mourillon (29 downregulated, 34 upregulated) (Supplementary Table 4; Table 2). Of these, 83 genes were annotated, and 6 were shared between the two comparisons (*CLEC1A, CRY2, HSPA5, IFI27, NFIL3* and *SCD*). GO term enrichment of the T0 DEGs further revealed four significantly over-represented pathways (FDR-adjusted p < 0.05). Three of these were associated with circadian rhythm regulation: *Phosphorylated BMAL1:CLOCK (ARNTL:CLOCK) activates expression of core clock genes* (p = 1.24E-8), *circadian clock* (p = 5.61E-6), and *CRY:PER:kinase complex represses transactivation by the BMAL1:CLOCK (ARNTL:CLOCK) complex* (p = 1.13E-3). The fourth enriched pathway was related to immunity (*modulation of host responses by IFN-stimulated genes*, p = 0.031). Eight KEGG pathways were significantly enriched based on these DEGs. These included *alpha-linolenic acid metabolism* (p = 0.003), *linoleic acid metabolism* (p = 0.004), *biosynthesis of unsaturated fatty acids* (p = 0.007), *ester lipid metabolism* (p = 0.014*), arachidonic acid metabolism* (p = 0.015), *fatty acid metabolism* (p = 0.023), *protein processing in the endoplasmic reticulum (ER)* (p = 0.025), and the *PPAR signaling pathway* (p = 0.032). Genes associated with the circadian rhythm (such as *CRY2* and *NFIL3*), ER stress and protein folding (*HSPA5*, *CAPN2* and *P58IPK*), and cell cycle pathways had lower transcriptional levels in Port, whereas genes related to immune responses (like *IFI27*), and lipid metabolism had predominantly higher transcriptional levels (Table 2). Although the detoxification pathways were not significantly enriched in the KEGG and GO terms of the DEGs, the *CYP1A1* gene had a higher transcriptional level in Port individuals compared to Méjean and Mourillon (Table 2).

**Table 2.**
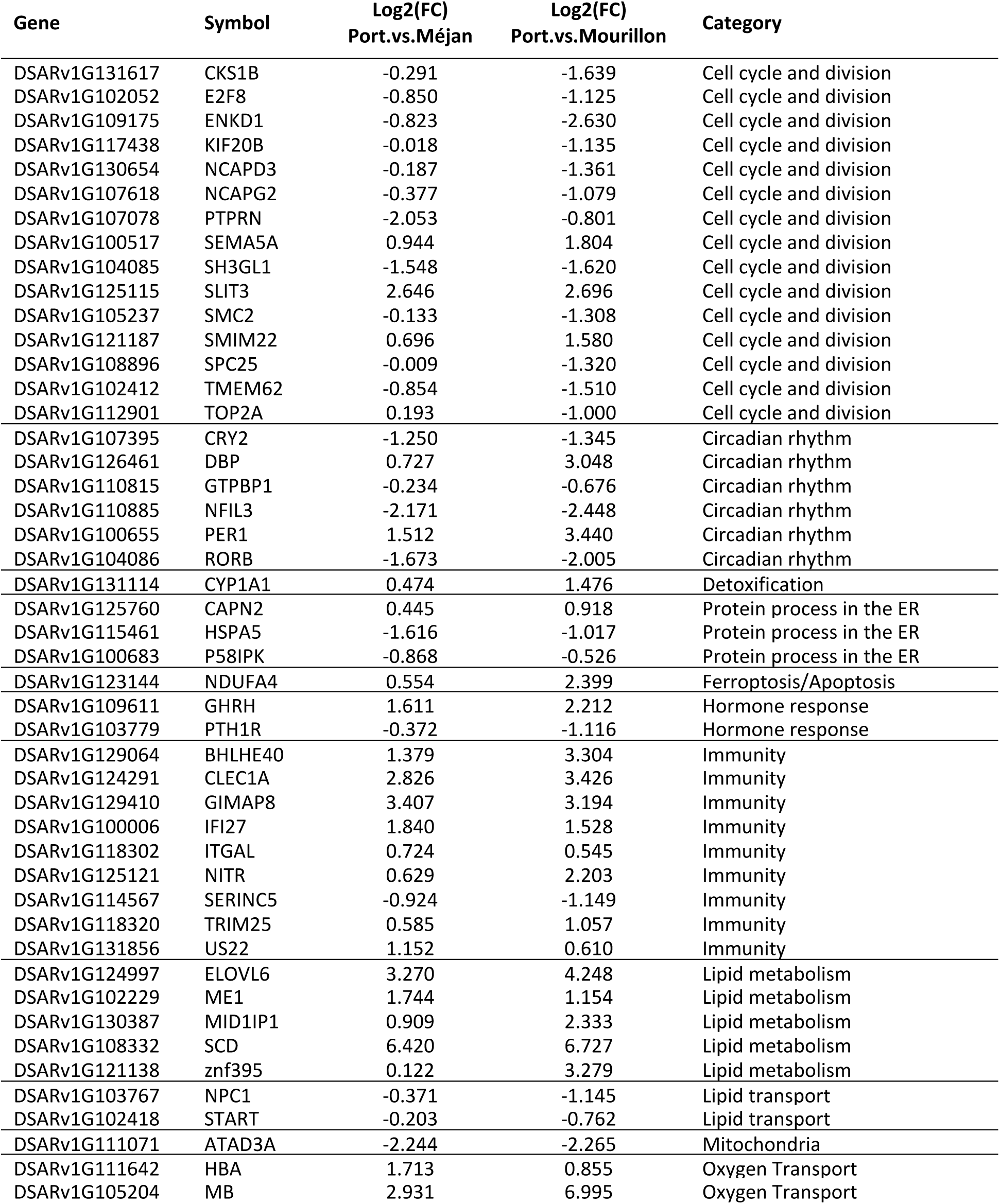
DEGs at T0 belonging to the different enriched KEGG and GO pathways, alongside their Log_2_(FC) in expression, sorted by functional category, for the Port versus Méjean comparison (reference= Méjean) and for the Port versus Mourillon comparison (reference= Mourillon).

#### DEA and functional enrichment between T0 and T1

Between T0 and T1, DEA identified 2693 DEGs in Méjean (1310 downregulated, 1383 upregulated), 3016 DEGs in Mourillon (1534 downregulated, 1482 upregulated), and 3733 DEGs in Port (1933 downregulated, 1800 upregulated) (Supplementary Table 5). Of all the DEGs identified, 938 genes were shared by the three populations and 322 genes were differentially expressed in Méjean and Mourillon (Port showing no significant change in the expression of those genes between T0 and T1). Interestingly, in the 322 DEGs, 5 genes were DEGs at T0 (two uncharacterized genes, ATAD3A, COL24A1 and CYP1A1; Figure 4). In addition to CYP1A1, we also identified three other CYP450 family genes among which CYP2J1 (Supplementary Table 5), for which the level of expression at T1 in Méjean and Mourillon matched that of Port at T0 and T1.

**Figure 4:**
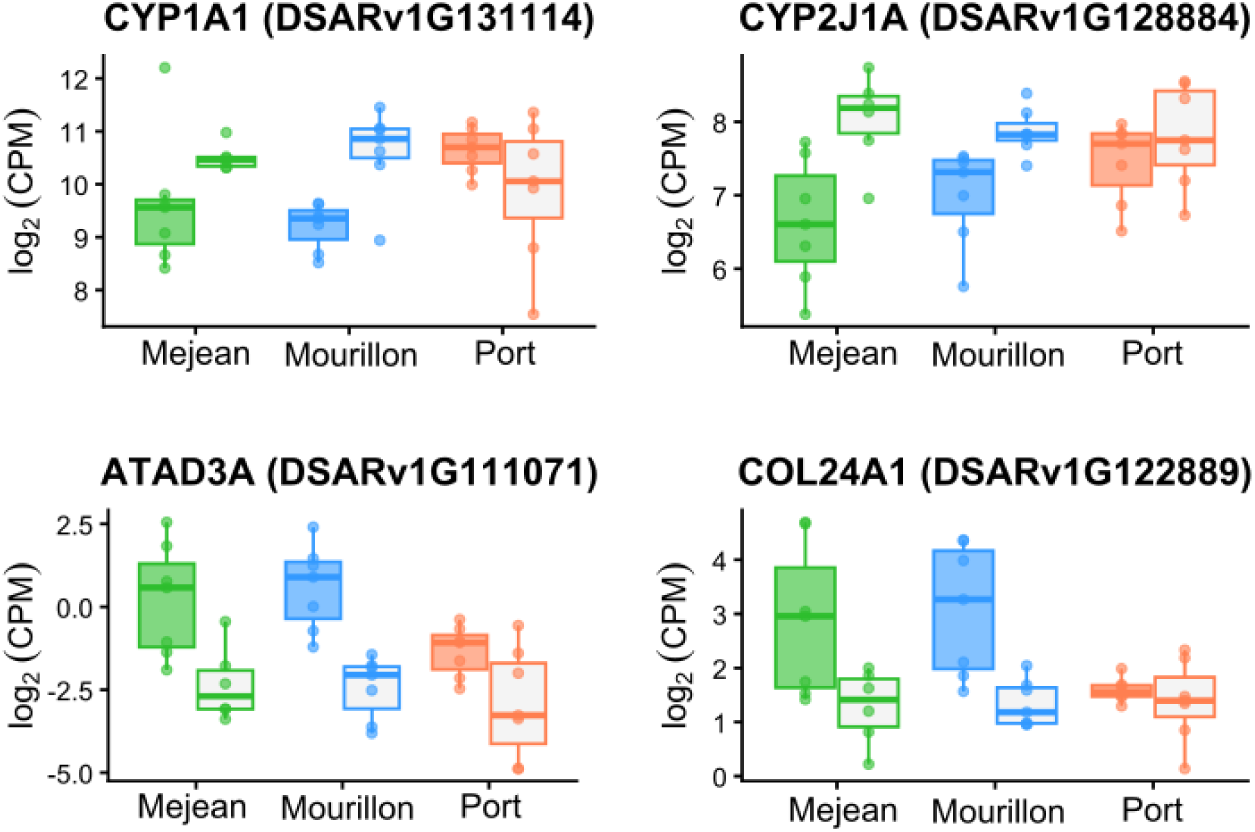
Gene expression level (log_2_(Count Per Million)) of key genes identified as DEGs between T0 and T1 in both Méjean and Mourillon, but not in the Port population.

Functional enrichment showed that 115, 50 and 78 GO terms were enriched for Méjean, Mourillon, and Port respectively (Supplementary Table 6), among which 18 were shared among the three populations, essentially belonging to cellular division and cellular processes (Table 3A). We found 55, 67 and 66 significantly enriched KEGG pathways for the Méjean, Mourillon and Port respectively (Supplementary Table 7), among which 33 were shared by the three populations (Table 3B). The 33 shared pathways showed a global coordinated response to caging characterized by detoxification, oxidative stress management, fatty acid and energy metabolism response, and metabolic reprogramming. Specifically, enriched pathways included the “*metabolism of xenobiotics by cytochrome P450”* and “*drug metabolism - other enzymes”,* the “*glycolysis / gluconeogenesis*”, “*insulin signaling pathway*”, “*adipocytokine signaling pathway*”, and several pathways linked to cell fate, such as *“apoptosis”, “necroptosis”, “autophagy”, or “cellular senescence”* (Table 3B). Two pathways were common with the T0 analysis: “*fatty acid metabolism”, “PPAR signaling pathway*”. We also identified 12, 11 and 14 population-specific pathways for Port, Méjean and Mourillon respectively (Supplementary Table 7). These pathways included among others the *“ribosome”, “ABC transporters”, “Glycerolipid metabolism”,* and *“protein processing in the ER”* for Port, *“spliceosome”, “Mismatch repair”, “RNA degradation”* and *“Mitophagy – animal”* for Méjean, and *“DNA repair”, “Tyrosine metabolism”, “Glycosphingolipid biosynthesis”* and *“Base excision repair”* for Mourillon (Supplementary Table 7).

**Table 3.**
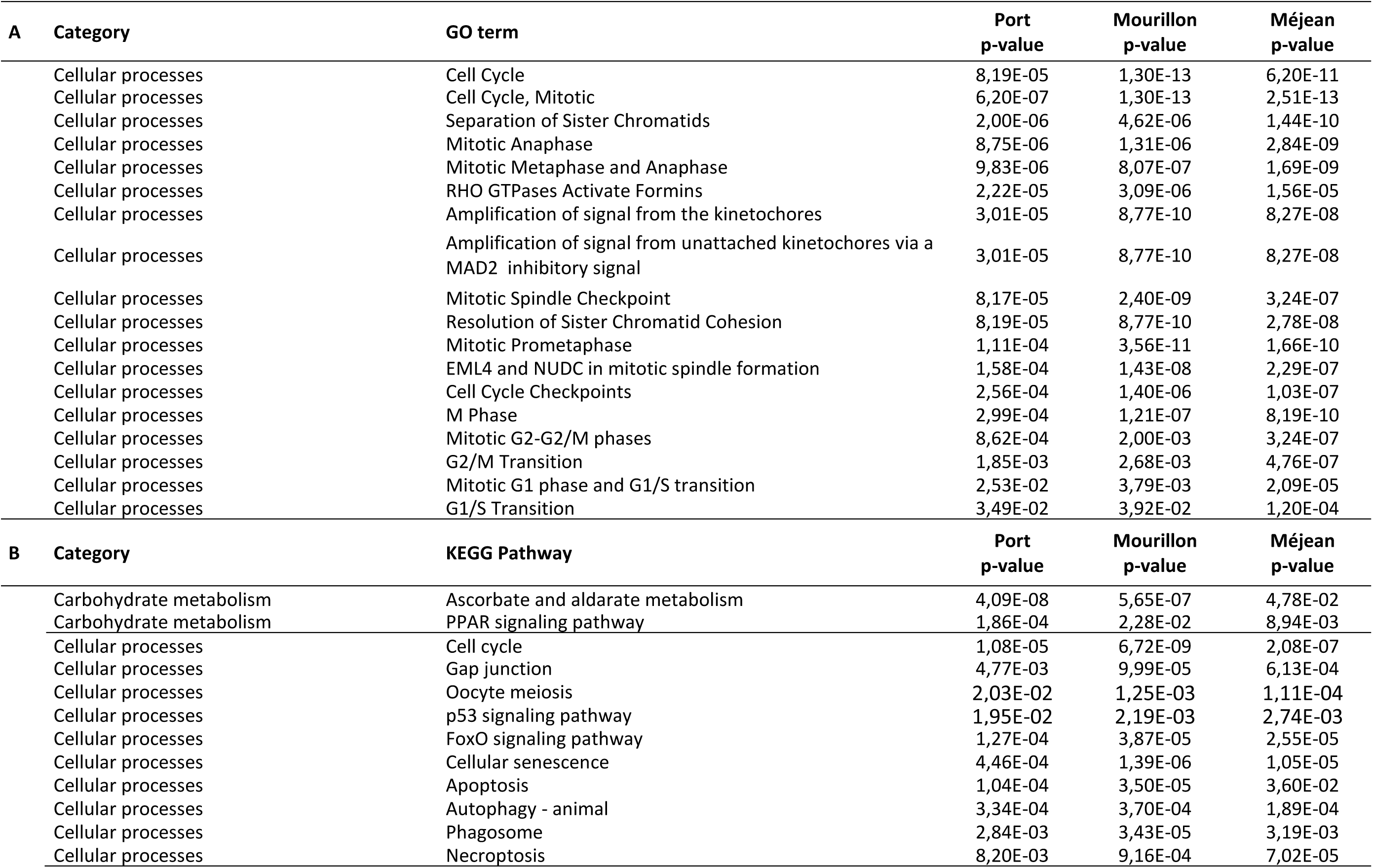
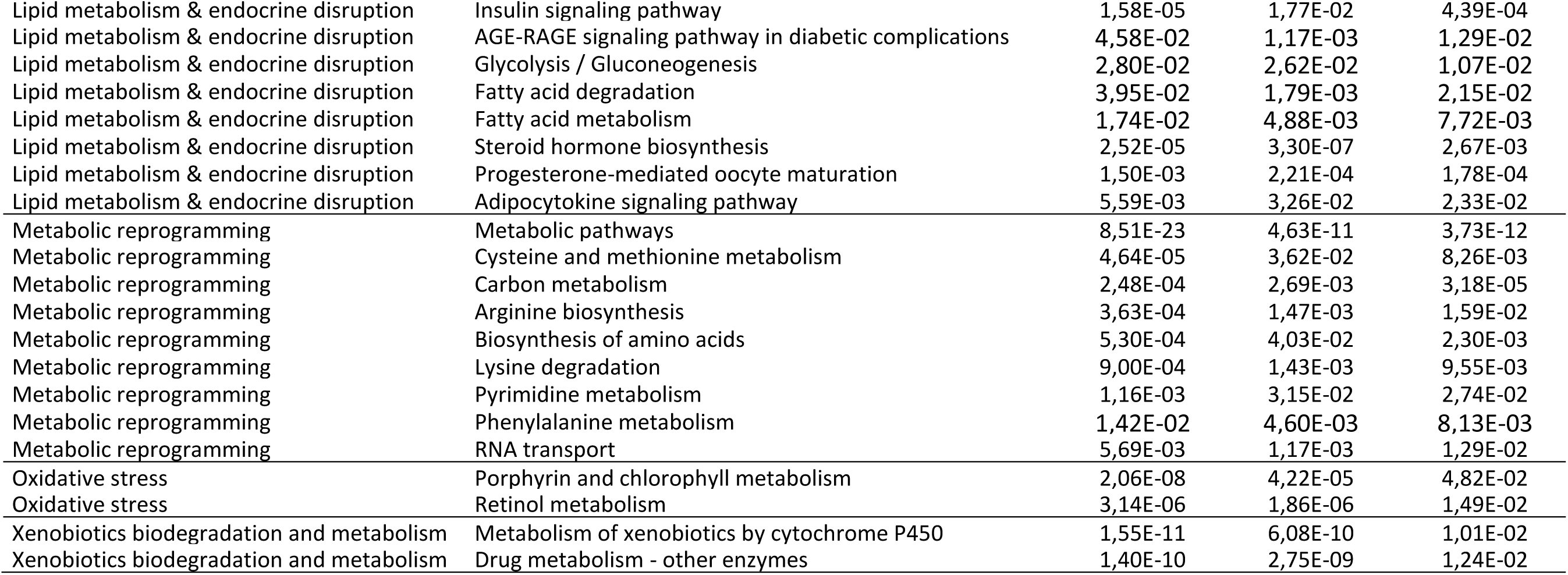
Enriched GO categories (A) and KEGG pathways (B) between T0 and T1 that are shared in Port, Mourillon and Méjean populations.

| A | Category | GO term | Port<br>p-value | Mourillon<br>p-value | Méjean<br>p-value |
| --- | --- | --- | --- | --- | --- |
|  | Cellular processes | Cell Cycle | 8,19E-05 | 1,30E-13 | 6,20E-11 |
|  | Cellular processes | Cell Cycle, Mitotic | 6,20E-07 | 1,30E-13 | 2,51E-13 |
|  | Cellular processes | Separation of Sister Chromatids | 2,00E-06 | 4,62E-06 | 1,44E-10 |
|  | Cellular processes | Mitotic Anaphase | 8,75E-06 | 1,31E-06 | 2,84E-09 |
|  | Cellular processes | Mitotic Metaphase and Anaphase | 9,83E-06 | 8,07E-07 | 1,69E-09 |
|  | Cellular processes | RHO GTPases Activate Formins | 2,22E-05 | 3,09E-06 | 1,56E-05 |
|  | Cellular processes | Amplification of signal from the kinetochores | 3,01E-05 | 8,77E-10 | 8,27E-08 |
|  | Cellular processes | Amplification of signal from unattached kinetochores via a<br>MAD2 inhibitory signal | 3,01E-05 | 8,77E-10 | 8,27E-08 |
|  | Cellular processes | Mitotic Spindle Checkpoint | 8,17E-05 | 2,40E-09 | 3,24E-07 |
|  | Cellular processes | Resolution of Sister Chromatid Cohesion | 8,19E-05 | 8,77E-10 | 2,78E-08 |
|  | Cellular processes | Mitotic Prometaphase | 1,11E-04 | 3,56E-11 | 1,66E-10 |
|  | Cellular processes | EML4 and NUDC in mitotic spindle formation | 1,58E-04 | 1,43E-08 | 2,29E-07 |
|  | Cellular processes | Cell Cycle Checkpoints | 2,56E-04 | 1,40E-06 | 1,03E-07 |
|  | Cellular processes | M Phase | 2,99E-04 | 1,21E-07 | 8,19E-10 |
|  | Cellular processes | Mitotic G2-G2/M phases | 8,62E-04 | 2,00E-03 | 3,24E-07 |
|  | Cellular processes | G2/M Transition | 1,85E-03 | 2,68E-03 | 4,76E-07 |
|  | Cellular processes | Mitotic G1 phase and G1/S transition | 2,53E-02 | 3,79E-03 | 2,09E-05 |
|  | Cellular processes | G1/S Transition | 3,49E-02 | 3,92E-02 | 1,20E-04 |
| B | Category | KEGG Pathway | Port<br>p-value | Mourillon<br>p-value | Méjean<br>p-value |
|  | Carbohydrate metabolism | Ascorbate and aldarate metabolism | 4,09E-08 | 5,65E-07 | 4,78E-02 |
|  | Carbohydrate metabolism | PPAR signaling pathway | 1,86E-04 | 2,28E-02 | 8,94E-03 |
|  | Cellular processes | Cell cycle | 1,08E-05 | 6,72E-09 | 2,08E-07 |
|  | Cellular processes | Gap junction | 4,77E-03 | 9,99E-05 | 6,13E-04 |
|  | Cellular processes | Oocyte meiosis | 2,03E-02 | 1,25E-03 | 1,11E-04 |
|  | Cellular processes | p53 signaling pathway | 1,95E-02 | 2,19E-03 | 2,74E-03 |
|  | Cellular processes | FoxO signaling pathway | 1,27E-04 | 3,87E-05 | 2,55E-05 |
|  | Cellular processes | Cellular senescence | 4,46E-04 | 1,39E-06 | 1,05E-05 |
|  | Cellular processes | Apoptosis | 1,04E-04 | 3,50E-05 | 3,60E-02 |
|  | Cellular processes | Autophagy - animal | 3,34E-04 | 3,70E-04 | 1,89E-04 |
|  | Cellular processes | Phagosome | 2,84E-03 | 3,43E-05 | 3,19E-03 |
|  | Cellular processes | Necroptosis | 8,20E-03 | 9,16E-04 | 7,02E-05 |

|  |  |  |  |  |
| --- | --- | --- | --- | --- |
| Lipid metabolism & endocrine disruption | Insulin signaling pathway | 1,58E-05 | 1,77E-02 | 4,39E-04 |
| Lipid metabolism & endocrine disruption | AGE-RAGE signaling pathway in diabetic complications | 4,58E-02 | 1,17E-03 | 1,29E-02 |
| Lipid metabolism & endocrine disruption | Glycolysis / Gluconeogenesis | 2,80E-02 | 2,62E-02 | 1,07E-02 |
| Lipid metabolism & endocrine disruption | Fatty acid degradation | 3,95E-02 | 1,79E-03 | 2,15E-02 |
| Lipid metabolism & endocrine disruption | Fatty acid metabolism | 1,74E-02 | 4,88E-03 | 7,72E-03 |
| Lipid metabolism & endocrine disruption | Steroid hormone biosynthesis | 2,52E-05 | 3,30E-07 | 2,67E-03 |
| Lipid metabolism & endocrine disruption | Progesterone-mediated oocyte maturation | 1,50E-03 | 2,21E-04 | 1,78E-04 |
| Lipid metabolism & endocrine disruption | Adipocytokine signaling pathway | 5,59E-03 | 3,26E-02 | 2,33E-02 |
| Metabolic reprogramming | Metabolic pathways | 8,51E-23 | 4,63E-11 | 3,73E-12 |
| Metabolic reprogramming | Cysteine and methionine metabolism | 4,64E-05 | 3,62E-02 | 8,26E-03 |
| Metabolic reprogramming | Carbon metabolism | 2,48E-04 | 2,69E-03 | 3,18E-05 |
| Metabolic reprogramming | Arginine biosynthesis | 3,63E-04 | 1,47E-03 | 1,59E-02 |
| Metabolic reprogramming | Biosynthesis of amino acids | 5,30E-04 | 4,03E-02 | 2,30E-03 |
| Metabolic reprogramming | Lysine degradation | 9,00E-04 | 1,43E-03 | 9,55E-03 |
| Metabolic reprogramming | Pyrimidine metabolism | 1,16E-03 | 3,15E-02 | 2,74E-02 |
| Metabolic reprogramming | Phenylalanine metabolism | 1,42E-02 | 4,60E-03 | 8,13E-03 |
| Metabolic reprogramming | RNA transport | 5,69E-03 | 1,17E-03 | 1,29E-02 |
| Oxidative stress | Porphyrin and chlorophyll metabolism | 2,06E-08 | 4,22E-05 | 4,82E-02 |
| Oxidative stress | Retinol metabolism | 3,14E-06 | 1,86E-06 | 1,49E-02 |
| Xenobiotics biodegradation and metabolism | Metabolism of xenobiotics by cytochrome P450 | 1,55E-11 | 6,08E-10 | 1,01E-02 |
| Xenobiotics biodegradation and metabolism | Drug metabolism - other enzymes | 1,40E-10 | 2,75E-09 | 1,24E-02 |

#### Differential expression analysis and functional enrichment at T1

DEA identified 500 differentially expressed genes (DEGs) between Port and Méjean (280 downregulated, 220 upregulated), and 51 DEGs between Port and Mourillon (34 downregulated, 17 upregulated) (Supplementary Table 8; Table 4). We identified 16 GO terms as enriched in the list of DEGs (Table 5), among which “*mitochondrial translation”*, “*mitochondrial ribosome-associated quality control”,* “*circadian clock”*, “*translation”*, “*heme signaling”* and “*plasma lipoprotein assembly”, remodeling, and clearance”*. A total of 25 KEGG pathways belonging to 13 broad categories were significantly enriched (Table 6), among which five pathways related to carbohydrate metabolism *(e.g. “glycolysis/gluconeogenesis”*, “*pentose phosphate pathway”*), four pathways related to lipid metabolism *(e.g. “fatty acid metabolism”*, “*glycerolipid metabolism*”), three pathways related to xenobiotics biodegradation and metabolism *(e.g. “drug metabolism - other enzymes”, “drug metabolism - cytochrome P450”*), three pathways related to genetic information processing (*e.g*. “*ribosome”*, “*RNA polymerase”*) (Table 6), and one pathway related to energy metabolism (“*oxidative phosphorylation* OXPHOS”).

**Table 4.** DEGs at T1 belonging to the different enriched KEGG and GO pathways, alongside their Log_2_(FC) in expression, sorted by functional category, for the Port versus Méjean comparison (reference= Méjean) and for the Port versus Mourillon comparison (reference= Mourillon).

| Gene | Symbol | Log2(FC)<br>Port.vs.Mej | Log2(FC)<br>Port.vs.Mour | Function |
| --- | --- | --- | --- | --- |
| DSARv1G103708 | YES1 | -1.196 | -0.609 | Cancer pathways |
| DSARv1G126597 | CYP2W1 | -1.436 | -0.764 | Cancer pathways |
| DSARv1G118077 | CYP2W1 | -1.054 | -0.873 | Cancer pathways |
| DSARv1G105234 | CLOCK | -1.315 | -1.735 | Circadian rhythm |
| DSARv1G115864 | CLOCK | -3.674 | -2.990 | Circadian rhythm |
| DSARv1G118739 | ARNTL | -2.314 | -1.527 | Circadian rhythm |
| DSARv1G118614 | ARNTL2 | -2.675 | -2.176 | Circadian rhythm |
| DSARv1G118615 | ARNTL2 | -2.773 | -2.084 | Circadian rhythm |
| DSARv1G107395 | CRY2 | -0.792 | -0.613 | Circadian rhythm |
| DSARv1G126461 | DBP | 3.190 | 2.346 | Circadian rhythm |
| DSARv1G104652 | NPAS2 | -1.407 | -0.793 | Circadian rhythm |
| DSARv1G113331 | NR1D1 | 2.452 | 1.690 | Circadian rhythm |
| DSARv1G100655 | PER1 | 3.529 | 3.041 | Circadian rhythm |
| DSARv1G132282 | PER3 | 1.566 | 1.201 | Circadian rhythm |
| DSARv1G104086 | RORB | -2.716 | -2.072 | Circadian rhythm |
| DSARv1G108774 | TBL1X | -1.165 | -0.740 | Circadian rhythm |
| DSARv1G116488 | ABCC3 | 0.710 | 0.654 | Drug metabolism - other enzymes |
| DSARv1G122467 | GST | 1.330 | 0.646 | Drug metabolism - other enzymes |
| DSARv1G117326 | MAOB | -1.526 | -0.937 | Drug metabolism - other enzymes |
| DSARv1G103263 | UGT1A1 | -1.601 | -0.853 | Drug metabolism - other enzymes |
| DSARv1G103262 | UGT1A1 | -1.356 | -0.682 | Drug metabolism - other enzymes |
| DSARv1G119314 | UGT2A3 | -1.045 | -0.436 | Drug metabolism - other enzymes |
| DSARv1G118631 | CES2 | -0.831 | -0.845 | Drug metabolism - other enzymes |
| DSARv1G124373 | CES5A | -0.835 | -0.562 | Drug metabolism - other enzymes |
| DSARv1G106040 | SULT1ST3 | -1.520 | -1.362 | Drug metabolism - other enzymes |
| DSARv1G123791 | TYMP | 1.099 | 0.751 | Drug metabolism - other enzymes |
| DSARv1G105914 | UCKL1 | 0.815 | 0.290 | Drug metabolism - other enzymes |
| DSARv1G115171 | HSP30 | -5.532 | -1.817 | Stress response |
| DSARv1G101445 | HSP90A | -3.076 | -1.819 | Stress response |
| DSARv1G113242 | HSPA1L | -3.398 | -1.728 | Stress response |
| DSARv1G120822 | ACADS | -0.921 | -0.513 | Lipid metabolism |
| DSARv1G107238 | ACOX1 | -1.018 | -0.633 | Lipid metabolism |
| DSARv1G100809 | AGK | 0.864 | 0.545 | Lipid metabolism |
| DSARv1G102780 | AP2S1 | -0.791 | -0.433 | Lipid metabolism |
| DSARv1G116048 | CPT2 | -1.336 | -0.989 | Lipid metabolism |
| DSARv1G119916 | DGKD | 0.769 | 0.189 | Lipid metabolism |
| DSARv1G111162 | ELOVL4 | -2.094 | -1.554 | Lipid metabolism |
| DSARv1G116479 | FASN | 1.594 | 0.658 | Lipid metabolism |
| DSARv1G110879 | GCDH | 1.376 | 0.873 | Lipid metabolism |
| DSARv1G117924 | GPAM | 1.566 | 0.592 | Lipid metabolism |
| DSARv1G114015 | MCAT | 0.940 | 0.381 | Lipid metabolism |
| DSARv1G108527 | MGLL | -1.113 | -0.847 | Lipid metabolism |
| DSARv1G102436 | NCEH1 | -2.462 | -1.775 | Lipid metabolism |
| DSARv1G102145 | PCSK6 | -1.080 | -0.672 | Lipid metabolism |
| DSARv1G111726 | PLPP3 | -0.888 | -0.433 | Lipid metabolism |
| DSARv1G100904 | SOAT2 | -2.031 | -1.415 | Lipid metabolism |
| DSARv1G113052 | ABCA1 | -1.270 | -0.596 | Lipid transport |
| DSARv1G122907 | ANGPTL3 | -2.495 | -1.343 | Lipid transport |
| DSARv1G103472 | ANGPTL4 | -1.340 | -0.946 | Lipid transport |
| DSARv1G102744 | APOA4 | -2.507 | -3.302 | Lipid transport |
| DSARv1G101135 | APOB | -2.109 | -2.055 | Lipid transport |
| DSARv1G107756 | APOF | -1.038 | -0.694 | Lipid transport |
| DSARv1G103767 | NPC1 | -1.456 | -0.996 | Lipid transport |
| DSARv1G109413 | MRPL1 | 0.787 | 0.286 | Mitochondrial processes and translation |
| DSARv1G113753 | MRPL13 | 0.571 | 0.564 | Mitochondrial processes and translation |
| DSARv1G106563 | MRPL14 | 1.178 | 0.374 | Mitochondrial processes and translation |
| DSARv1G106492 | MRPL2 | 0.779 | 0.382 | Mitochondrial processes and translation |
| DSARv1G127832 | MRPL21 | 0.935 | 0.356 | Mitochondrial processes and translation |
| DSARv1G131323 | MRPL3 | 0.800 | 0.221 | Mitochondrial processes and translation |
| DSARv1G113540 | MRPL39 | 1.083 | 0.531 | Mitochondrial processes and translation |
| DSARv1G108663 | MRPL42 | 0.976 | 0.425 | Mitochondrial processes and translation |
| DSARv1G127829 | MRPL45 | 0.714 | 0.218 | Mitochondrial processes and translation |
| DSARv1G112068 | MRPL51 | 0.868 | 0.040 | Mitochondrial processes and translation |
| DSARv1G120984 | MRPL52 | 0.971 | 0.339 | Mitochondrial processes and translation |
| DSARv1G124689 | MRPS11 | 0.876 | 0.238 | Mitochondrial processes and translation |
| DSARv1G109682 | MRPS16 | 0.674 | 0.674 | Mitochondrial processes and translation |
| DSARv1G122817 | MRPS17 | 0.805 | 0.171 | Mitochondrial processes and translation |
| DSARv1G101383 | MRPS18A | 0.694 | 0.187 | Mitochondrial processes and translation |
| DSARv1G128012 | MRPS18B | 0.699 | 0.201 | Mitochondrial processes and translation |
| DSARv1G117029 | MRPS26 | 0.899 | 0.434 | Mitochondrial processes and translation |
| DSARv1G110062 | MRPS27 | -2.048 | -1.704 | Mitochondrial processes and translation |
| DSARv1G109263 | MRPS30 | 0.815 | 0.248 | Mitochondrial processes and translation |
| DSARv1G130859 | MRPS5 | 0.635 | 0.291 | Mitochondrial processes and translation |
| DSARv1G110328 | MRPS9 | 0.788 | 0.359 | Mitochondrial processes and translation |
| DSARv1G103441 | RPL5 | 1.154 | 0.427 | Mitochondrial processes and translation |
| DSARv1G114182 | RPS2 | 0.914 | 0.345 | Mitochondrial processes and translation |
| DSARv1G124794 | TUFM | 0.921 | 0.318 | Mitochondrial processes and translation |
| DSARv1G112526 | GADD45B | 1.523 | 1.481 | Oxidative stress |
| DSARv1G107118 | GPX4 | -1.296 | -0.890 | Oxidative stress |
| DSARv1G112258 | CROT | -1.258 | -0.693 | Peroxisome |
| DSARv1G110605 | PHYH | -1.264 | -1.121 | Peroxisome |
| DSARv1G123002 | PIPOX | -1.550 | -0.993 | Peroxisome |
| DSARv1G114830 | SLC27A2 | -1.557 | -1.220 | Peroxisome |

**Table 5.** Enriched KEGG pathways at T1 identified in at least one comparison. Significance was set at p-value<0.05.

| Category | #Term | p-Value |
| --- | --- | --- |
| Amino acid metabolism | Glycine, serine and threonine metabolism | 5.99E-03 |
| Amino acid metabolism | Biosynthesis of amino acids | 1.59E-02 |
| Carbohydrate metabolism | Citrate cycle (TCA cycle) | 2.29E-04 |
| Carbohydrate metabolism | Ascorbate and aldarate metabolism | 1.47E-02 |
| Carbohydrate metabolism | Pentose and glucuronate interconversions | 2.78E-02 |
| Carbohydrate metabolism | Glycolysis / Gluconeogenesis | 3.17E-02 |
| Carbohydrate metabolism | Pentose phosphate pathway | 3.73E-02 |
| Carbon metabolism | Carbon metabolism | 2.26E-05 |
| Energy metabolism | Oxidative phosphorylation | 3.32E-04 |
| Genetic Information Processing | Ribosome | 2.32E-07 |
| Genetic Information Processing | Ribosome biogenesis in eukaryotes | 5.02E-03 |
| Genetic Information Processing | RNA polymerase | 2.78E-02 |
| Lipid metabolism | Fatty acid metabolism | 5.02E-03 |
| Lipid metabolism | Fatty acid degradation | 2.13E-02 |
| Lipid metabolism | Steroid hormone biosynthesis | 2.56E-02 |
| Lipid metabolism | Glycerolipid metabolism | 2.66E-02 |
| Lipid oxidation | Peroxisome | 4.85E-02 |
| Lipid oxidation | PPAR signaling pathway | 2.97E-03 |
| Metabolic pathways | Metabolic pathways | 1.20E-07 |
| Metabolism of other amino acids | beta-Alanine metabolism | 4.26E-02 |
| Oxidative stress | Porphyrin and chlorophyll metabolism | 4.26E-02 |
| Oxocarboxylic acid metabolism | 2-Oxocarboxylic acid metabolism | 1.79E-02 |
| Xenobiotics biodegradation and metabolism | Drug metabolism - other enzymes | 5.38E-04 |
| Xenobiotics biodegradation and metabolism | Drug metabolism - cytochrome P450 | 9.67E-03 |
| Xenobiotics biodegradation and metabolism | Metabolism of xenobiotics by cytochrome P450 | 4.54E-02 |

**Table 6.** Enriched GO categories for the T1 DEA analyses, with their enrichment ratio and associated p-value, with an FDR correction of 5%. Significance was set at p-value < 0.05.

| Category | Pathway name | p-value |
| --- | --- | --- |
| Circadian rhythm | Circadian clock | 7.42E-05 |
| Circadian rhythm | Phosphorylated BMAL1:CLOCK (ARNTL:CLOCK) activates expression of core clock genes | 1.04E-08 |
| Circadian rhythm | The CRY:PER:kinase complex represses transactivation by the BMAL:CLOCK | 1.22E-06 |
| Circadian rhythm | Phosphorylation of CLOCK, acetylation of BMAL1 (ARNTL) at target gene promoters | 2.16E-03 |
| Circadian rhythm | Expression of BMAL (ARNTL), CLOCK, and NPAS2 | 4.93E-03 |
| Cell cycle | Attenuation phase | 2.39E-02 |
| Genetic information process | Translation | 1.75E-03 |
| Genetic information process, mitochondria | Mitochondrial translation | 2.27E-06 |
| Genetic information process, mitochondria | Mitochondrial translation elongation | 1.04E-08 |
| Genetic information process, mitochondria | Mitochondrial translation initiation | 4.44E-08 |
| Genetic information process, mitochondria | Mitochondrial ribosome-associated quality control | 7.62E-08 |
| Genetic information process, mitochondria | Mitochondrial translation termination | 1.80E-07 |
| Lipid metabolism | NR1H2 and NR1H3-mediated signaling | 6.14E-03 |
| Lipid metabolism | Plasma lipoprotein assembly, remodeling, and clearance | 2.44E-02 |
| Signaling | Heme signaling | 2.09E-02 |
| Signaling | Signaling by Nuclear Receptors | 2.34E-02 |

When examining individual genes, DEGs contained more genes involved with circadian rhythm compared to DEGs at T0, among which two *CLOCK,* one *PER3*, and two *ARNTL2*. Most of these genes were downregulated in Port population compared to the two other sites (Table 4). Also, while *CYP1A1* has a similar expression in all three populations at T1, many genes involved in phase II (*e.g.*, two *UGT1A1*, one *GST,* one *SULT1ST3*) and phase III (*ABCC3*) of the xenobiotic detoxification mechanisms are downregulated in individuals originating from Port compared to Méjean and Mourillon (Table 4). Stress-related genes (*HSP30*, *HSPA1L* and *HSP90A*), lipid metabolism genes (*APOA4*, *APOB*, *ELOVL4* among others) and peroxisome related genes (*CROT*, *PHYH*, *PIPOX* and *SLC27A2*) are also less expressed in Port. Among the lipid-related genes, *NPC1*, involved in lipoprotein-derived cholesterol transport, was already expressed at lower levels in Port at T0 (Table 2), and this reduced expression persisted at T1 (Table 4). In addition, several *MRPL* and *MRPS* genes, involved in energy metabolism and OXPHOS pathways are upregulated in Port (Table 4). Finally, two genes involved in cancer pathways (*CYP2W1* and *YES1*) are upregulated in Méjean and Mourillon compared to Port (Table 4).

## Discussion

Coastal regions are experiencing increasing urbanization, most of which is due to port constructions that leads to substantial modifications in habitat structure and environmental conditions [10]. These changes can significantly impact the organisms that inhabit these areas, and especially coastal species juveniles that use them as nurseries. However, the specific effects of port environments on the physiology and health of juvenile fish remain poorly understood, particularly for species that utilize these habitats on a transient basis, during early life stages.

Our findings demonstrate that juvenile *D. vulgaris* migrating into port habitats for growth are exposed to a complex and stressful environment that significantly alters their physiology and transcriptomic response. While our results could suggest that these individuals possess the capacity for short-term acclimation to such conditions, the potential long-term costs of these physiological and molecular response remain uncertain and merit further investigation.

### Genetic structuring among sampled sites

It is well documented that gene expression can be modulated both by the environment and genetic differences [56,57]. RAD-sequencing identified a low but significant genetic structuring among the three studied sites, with Méjean being differentiated from Port and Mourillon, and its individuals showing a higher genetic diversity. However, multivariate methods (PCA) and unsupervised hierarchical clustering failed to identify more than one genetic cluster. A few individuals from Méjean could thus originate from a slightly different reproductive pool of adults, while all the other individuals of our study originated from a common reproductive. *Diplodus* species in the Mediterranean Sea commonly show weak to non-existent genetic structuring across their geographic range, suggesting a global high gene flow and capacity for long distance dispersal [58–61]. The small genetic divergence we observe here could have a small influence on the differences in response to port conditions between Méjean and the other two populations. However, this influence should remain small, and the influence of the environment (port conditions and caging response) is a more likely factor for their variation in response.

### Impact of port conditions on fish physiology: bigger and fatter fish

Biometric analysis of T0 juveniles showed that Port fish exhibited an overall better body condition (heavier and longer) than those of Méjean and Mourillon, and had more triglycerides in their muscle tissues. This difference could be linked to nutrient enrichment commonly observed in port environments, directly exposed to anthropogenic urban and industrial runoff that can significantly elevate nitrogen and phosphorus levels [3,62]. Such enrichment can stimulate primary productivity and could increase food availability for *D. vulgaris* juveniles, which feed primarily on algae, copepods, and amphipods [63], although we did not directly measure these water parameters here. Nutrient enrichment could also explain the significant growth of the Méjean individuals between T0 and T1, Méjean being the geographically furthest from Port. Additionally, the higher expression of two genes involved in oxygen transport (*MB* and *HBA*) in Port at T0, may indicate more frequent episodes of hypoxia (a decrease in oxygen availability due to enhanced primary production), as commonly reported in port environments that are more sheltered than open coastal sea [64].

Another factor that may account for the greater size and lipid reserves observed in Port individuals at T0 is reduced predation pressure, which could allow them to forage more extensively and for longer durations. Indeed, predation pressure has been reported to be lower in artificial fish nurseries (AFNs) located within port environments compared to those outside [65]. Finally, the presence of pollutants themselves could explain these differences. Indeed, organic contaminants and metals, and this accumulation can directly influence the metabolism of fatty acids in living organisms [66], leading to a possible increase in total lipids, triglycerides and total cholesterol [67,68] and an alteration of molecular pathways such as PPAR, oxidation pathways, gluconeogenesis, and lipid metabolism [68]. The fact that all these pathways were found differentially regulated in the non-port population were confronted to port environment (*i.e.* after caging) could support this hypothesis.

### Differential regulation of detoxification, lipid metabolism and circadian rhythm pathways in Port at T0

EROD activity was significantly elevated in Port at T0, and the expression of *CYP1A1*, the gene that induces EROD activity, was also significantly higher in Port individuals at T0. The liver plays a key role in xenobiotic detoxification, a process that unfolds in three main stages: Phase I involves the modification of contaminants through oxidation, typically by cytochrome P450 enzymes, often rendering them more reactive [69]. During Phase II, these modified compounds are conjugated with polar molecules (*e.g.*, glucuronic acid, glutathione) by enzymes such as UDP-glucuronosyltransferases (*UGTs*), sulfotransferases (*SULTS*), N-acetyltransferases (NATs) to enhance solubility and facilitate elimination [70]. Finally Phase III entails the export of these conjugated products by membrane transporters such as solute carrier (SLC) and ATP binding cassette (ABC) proteins, allowing their excretion via bile or urine [71]. Detoxification through phase I enzymes, particularly cytochrome P450s, is known to generate reactive intermediates that can lead to oxidative stress and endoplasmic reticulum (ER) stress. DEGs at T0 in Port were significantly enriched for functions related to protein folding, with many of these genes showing lower transcriptional levels in Port, a pattern that may contribute to prolonged ER stress and indicate compromised homeostasis. Immune pathways were also enriched among DEGs, notably with the elevated expression of several interferon-stimulated genes (ISGs), including *IFI27* and *TRIM25*. ISGs provide frontline defense against RNA and DNA viruses, as well as intracellular pathogens, and also play key roles in modulating adaptive immunity [72,73]. Their higher expression level in Port indicates a possible increased pathogenic pressure from the port environment.

Accordingly with the biometric differences observed between Port individuals and Méjean and Mourillon at T0, transcriptomic analysis of liver tissue from Port at T0 showed elevated transcriptional levels of several genes involved in lipid metabolism and transport, which could affect lipid accumulation in muscle. Indeed, xenobiotics are known to disrupt lipid metabolism in fish. For instance, increased hepatic lipid storage associated with contaminated diets has been documented in another Sparidae, *Sparus aurata* [74], and experimental exposure to contaminated water has been shown to impair hepatic lipid metabolism in goldfish [75]. In addition, PPAR signaling and lipid homeostasis can be impaired by the presence of PCBs, as demonstrated in the killifish [76]. Interestingly, we also found that transcript levels of *NPC1*, a gene that mediates the transport of lipoprotein-derived cholesterol from late endosomes and lysosomes to other cellular compartments, was lower in Port. *NPC1* plays a central role in maintaining intracellular cholesterol homeostasis, and its reduced expression has been associated with early-onset and severe obesity in both humans and mice [77].

Pathways related to the circadian rhythm were also enriched in the DEGs list, with most of the genes exhibiting a lower expression in individuals from Port, while *PER1* and *DBP* had a higher expression. *PER1* is a circadian oscillator which, through its accumulation in the nucleus, participates in the inhibition of the transcriptional activity of CLOCK:BMAL1 complex. Its downregulation and subsequent degradation cancel the suppression of BMAL1:CLOCK activity, allowing the initialization of a new oscillatory cycle [78]. *PER1* expression is light-sensitive and has been shown to increase in the liver of rodents exposed to artificial light at night (ALAN), particularly blue wavelengths [79,80]. Global estimates from satellite imagery suggest that approximately 22.2% of the world’s coastlines were affected by ALAN as of 2010, with European coastlines among the most impacted (53.4%) [81]. Coastal light pollution is increasing at a rate of ∼6% annually and is known to disrupt a range of biological processes in marine organisms, including behavior, orientation, reproduction, and overall physiological function (reviewed in [81]). Elevated expression of *PER1* and *DBP* may thus reflect an adaptive or stress-induced response to chronic light exposure in port environments. Additionally, genes involved in circadian rhythm are also known to predispose to the development of certain type of cancers [82], and are often downregulated in tumors, such as in melanomas in the zebrafish [83].

### Response to caging in port conditions

To investigate the physiological and molecular response of fish juveniles to port conditions, we caged individuals from all three populations in port conditions for one month and monitored their physiological and molecular response. We expected that the “naive” populations, Méjean and Mourillon, would show a difference in growth, weight and/or lipid reserves, and gene expression differences for detoxification related pathways as well as stress-related pathways (also due to caging), while Port would mainly show a difference in stress related pathways.

After one month of caging, we observed an absence of growth in Mourillon and Port individuals, while Méjean individuals slightly grew in length. We also observed a sharp decline in muscle lipid reserves in all three populations. We believe that, despite the installation of colonized oyster shell reefs, the caging conditions did not provide sufficient food resources over the long term to sustain juvenile growth. This may have influenced some of the results described below, by contributing to the activation of some molecular pathways. It could also potentially explain why Port individuals seemed to better cope with the constraints of the caging conditions (starvation and pollution) and modulated their physiological responses relative to the other populations, since they had greater lipid reserves. However, no mortality or signs of cannibalistic behavior (*e.g.* missing caudal fins or tegument injuries) were observed in the cages after one month. Moreover, Méjean individuals managed to grow, which could indicate that our results are still relevant. Nevertheless, caging conditions should be improved in future experiments, for example by reducing the exposure duration or by improving food substrates in the cage. Indeed, a similar experimental design using a shorter 15-day caging period has been successfully applied in sea bass and turbot without affecting growth or body condition [17].

EROD activity strongly diminished in Port individuals while it significantly increased in Méjean individuals (the site the furthest from Port), and remained stable in Mourillon individuals. This contrasting pattern does not entirely correlate with transcriptomic levels of CYP1A1, which significantly increased in Méjean and Mourillon individuals, and remained stable in Port individuals. The reduced EROD activity despite higher levels of CYP1A1 could directly be due to starvation of the Port population during caging which is known to reduce EROD activity despite the presence of contaminants [84].

Between T0 and T1, all populations modified the level of expression of genes involved in response to xenobiotic pathways, including cytochrome P450 and associated enzymatic systems, and antioxidant-related pathways. In all three populations, enrichment of core metabolic pathways could suggest a reallocation of energetic resources toward maintenance and repair functions. The enrichment of lipid metabolism and endocrine-related pathways, including PPAR signaling and steroid hormone biosynthesis could imply a potential metabolic and hormonal disruption. At the cellular level, the simultaneous involvement of apoptosis, autophagy, necroptosis, and senescence pathways in all the populations highlights ongoing regulation of cell fate. Hence, all three populations showed a very similar response to caging, including the differential regulation of detoxification pathways, which was not expected for Port. Of the 322 genes identified as differentially expressed genes (DEGs) between T0 and T1 at Méjean and Mourillon, only 5 overlapped with the DEGs of Port at T0. They belonged to the detoxification pathway (CYP1A1), mitochondrial membrane (ATAD3), collagen and integrin pathway (COL24A), while two genes remain unannotated. This indicates that the response to caging does not entirely align with the differences we identified at T0. It should be noted, however, that our results à T1 are not only influenced by port conditions but also by the caging effect, particularly the food deprivation experienced by the fish at the end of the month of caging. Specifically, fasting attenuated glucose and lipid anabolism, and enhanced catabolic pathways, fatty acid transport, PPAR signaling and liver antioxidant capacity in the javelin goby [85]. Hence, our results should be taken carefully and not be interpreted exclusively from the perspective of port conditions.

### Acclimation to port conditions

Acclimation enables organisms to adjust to new or changing environmental conditions within their lifetime. This process is underpinned by phenotypic plasticity, whereby a single genotype can produce a range of phenotypes in response to environmental variability [86]. Through such physiological adjustments, phenotypic plasticity offers a mechanism for short-term resilience to environmental stressors, including those associated with anthropogenic pressures. In our study, we aimed to test for the presence of acclimation in Port individuals. While when comparing population specific response to caging (T0 versus T1), we did not find a significant difference in the amplitude (number of genes) and the type of molecular response (enriched pathways) among populations, the presence of an acclimation of Port individuals could also result in a difference in expression of certain genes after caging (T1), possibly associated to differences in baseline expression. Our results show that pathways involved in detoxification, oxidative stress, lipid metabolism, oxidation and transport, and energy-related pathways were differentially regulated between Port and Méjean and Mourillon. While this might reflect the presence of acclimation in Port individuals, our results should also here be carefully interpreted, because of the likely influence of starvation.

Regarding the response to xenobiotics, although *CYP1A1* expression did not differ among populations after one month of caging, several other genes involved in detoxification pathways, such as *UGT1A1, GST* and *SULT1ST3*, were more expressed in Méjean and Mourillon compared to Port, potentially indicating enhanced detoxification activity in these individuals. Stress-response genes also exhibited higher transcriptional levels in Méjean and Mourillon, including *HSP30, HSP90A,* and *GADD45B,* the latter being known to increase its expression during hepatic oxidative stress [87]. In Méjean in particular, hepatic oxidative stress may be linked to the elevated EROD activity observed at T1, together with the increased expression of *CYP1A1* relative to pre-caging conditions. Concurrently, several genes involved in lipid β-oxidation, such as *ACADS*, *ACOX1*, and *CPT2*, were more expressed in Méjean and Mourillon than in Port at T1, which may have further contributed to increased oxidative stress.

The lower expression of lipid transport-related genes observed in Port at T1 may reflect improved energy conservation under caging conditions compared with Méjean and Mourillon. In addition, the higher expression of several nuclear-encoded mitochondrial ribosomal proteins, which contribute to mitochondrial protein synthesis and the assembly of oxidative phosphorylation (OXPHOS) complexes, may indicate a greater capacity to maintain efficient energy production in Port individuals. This response could represent an acclimation strategy aimed at sustaining ATP production under challenging environmental conditions. Alternatively, Port individuals may have been able to maintain OXPHOS pathway activation for longer periods due to their higher triglyceride reserves prior to caging, reserves that remained greater than those of the other two populations even after one month of caging.

Of greater concern, the Méjean and Mourillon populations had higher expression levels of several genes associated with oncogenic processes at T1, compared to Port. Among these, *CYP2W1* is characterized by strong tumor-specific expression and limited activity in normal tissues [88], while *YES1* is a well-characterized proto-oncogene [89]. In addition, many genes of the circadian rhythm were also differentially regulated between Port and Méjean and Mourillon, including *ARNTL* and two *ARNTL2* transcripts, which showed a lower transcript level in the Port. *ARNTL2* is a putative oncogene implicated in tumorigenesis and the regulation of cancer-associated immune responses, with elevated expression being associated with an immunosuppressive tumor microenvironment [90]. Also, *PER3* had a much lower expression in Méjean and Mourillon at T1 compared to the port, which is also a phenomenon observed in melanoma cells in zebrafish [83]. The contrasted expression level of these genes in Méjean and Mourillon individuals may suggest an increased susceptibility to cancer-related processes under port environmental conditions compared with port-originating individuals.

## Conclusions and perspectives

This study provides evidence that juvenile fish utilizing port environments as nursery grounds are exposed to complex and chronic stressors that significantly alter their physiology and molecular responses. Individuals originating from the polluted port seemed to exhibited transcriptional signatures consistent with acclimation, including reduced activation of detoxification pathways and increased expression of energy metabolism genes, particularly those related to mitochondrial function. In contrast, individuals who spent their growth phase outside the port showed pronounced activation of detoxification and immune pathways following one month of caging, including the upregulation of genes implicated in oncogenesis, raising questions about the long-term health consequences of early-life exposure to contaminated habitats. Aquatic pollution can lead to increased risk of cancerous growth and neoplasia in many species, and significantly affect fish health [91,92], even in later generations. Putative transgenerational effect of early-life exposure to pollutant is of particular concern for coastal fish species using ports as nurseries, since it can cause long-lasting behavioral, developmental and skeletal defects, persisting across multiple generations even without further exposure [93,94].

Our results are of particular relevance regarding the recent efforts to rehabilitate the nursery function in marine ports. Typical restoration efforts include deploying 3D artificial structures on the concrete walls of docks, under floating pontoons, or deposited directly at the bottom, to act as a reef to enhance habitat complexity and provide food resources and shelter from predators. Although some studies have reported successful recolonization of these artificial structures at small scale [95–99], their long-term ecological effectiveness remains uncertain [97,100], and these rehabilitation strategies primarily address the physical structure of habitats and often overlook the importance of density-independent factors, such as the quality of the surrounding environment. If the conditions in rehabilitated sites negatively affect survival, growth, or reproduction, these structures risk functioning as *ecological traps* [101] and the intended benefits of rehabilitation could be compromised, ultimately hindering population recovery and conservation goals. This highlights the importance of assessing not only patterns of habitat use but also the fitness consequences associated with occupation of restored environments.

## Acknowledgements

We thank Benoist de Vogüe, Fabienne Chavanon, Christophe Ravel, Leelou Chouteau, Elise Georges, Nicolas Martin and Etienne Joubert for their precious contribution to field sampling and fish dissections.

## Authors contributions

CMOR, EF, EBB, KM and MB designed the study and secured funding. CMOR, EF and MB sampled the fish. EMM and FC performed DNA and RNA extractions. EF and CMOR performed EROD analysis. CMOR and MR analyzed all the data. All authors contributed to the writing and editing of the manuscript. A CC-BY public copyright license has been applied by the authors to the present document and will be applied to all subsequent versions up to the Author Accepted Manuscript arising from this submission, in accordance with the grant’s open access conditions.

## Ethics approval statement

Fish were captured in accordance with French regulations on wildlife sampling, under scientific fishing authorization DIRM/112 of March 16, 2023 delivered by the Direction Interrégionale de la Mer Méditerrannée (DIRMM).

All experimental procedures involving animals were conducted in accordance with French and European regulations on animal welfare and experimentation (Directive 2010/63/EU). The study received ethical approval from the French Ministry of Higher Education and Research (MESR) under the authorization number APAFIS #41103-2023022212032882 v4, issued on the 16^th^ June 2023.

## Data availability statement

All the data (physiological measurements, VCF file for RAD-seq and raw genes counts for RNA-seq) alongside their statistical scripts are available at the gitlab repository https://gitlab.ifremer.fr/cr7a4b0/diplodus_toulon.git. Raw data for RNA-seq and RAD-seq are available from the ENA database (**Accession Number accessible upon acceptance of the article**), and the bioinformatic scripts to obtain the VCF file and raw gene counts are available in the gitlab repository cited above.

## Supplementary Material

Supplementary Table 1. Biometric measurement of *D. vulgaris* individuals from T0 and T1

Supplementary Table 2. Muscle lipid content results for individuals at T0 and T1

Supplementary Table 4: List of DEGs at T0 for each DEA comparison.

Supplementary Table 5: List of DEGs between T0 and T1 for each DEA comparison.

Supplementary Table 6: List of enriched GO terms between T0 and T1.

Supplementary Table 7: KEGG term enrichment results between T0 and T1 from KOBAS-i.

Supplementary Table 8: List of DEGs at T1 for each DEA comparison.

